# Synthesis and characterization of NIR fluorescent BODIPY-Xanthene hybrid dyes

**DOI:** 10.64898/2026.10.04.756502

**Authors:** Georgia Nigianni, Sotirios Katsamakas, Eleni Alexandratou, Kyriakos C. Prousis

## Abstract

The development of NIR fluorescent dyes with favorable optical properties is of great importance especially for applications within the field of optical imaging. In this study, we report the design, synthesis and photophysical characterization of novel BODIPY-Xanthene hybrid dyes obtained via Knoevenagel condensation between an asymmetrical BODIPY core and the functionalized xanthene aldehydes intermediates. The synthesized dyes exhibit absorption (667 nm) and emission bands (720–722 nm) within the NIR region. Detailed UV-Vis and fluorescence studies were performed and molar extinction coefficients and stokes shifts also calculated. Moreover, findings highlighting the influence of structural modifications on the optical behavior of the hybrid dyes and their potential utility in optical imaging applications are reported.

## 1. Introduction

Organic fluorophores have gained considerable attention the recent years since they are widely used as molecular tools for optical imaging and biomedical applications. Most of the existing fluorescent dyes, absorb and emit light within the visible region (between 400 to 650 nm), showcase several limitations including tissue autofluorescence and significant drawbacks such as the low tissue penetration due to the ability of biological tissues to absorb in the same region. Improved optical imaging has been achieved using fluorophores emitting light within the NIR window (650 nm to 900 nm) and beyond. The advantages of NIR fluorescent dyes include minimal interfering absorption and fluorescence from biological samples, reduced scattering and enhanced tissue penetration depth. [1] Interestingly, targeted NIR fluorescent probes enable emerging image guided surgical interventions, providing real-time definition of tumour margins. [2] Currently, there is an increasing demand for the development of photostable NIR dyes presenting large Stokes shifts, due to the emerging interest in optical *in vivo* imaging as well as for their applications in the modification of nanomaterials.

There are several important classes of well-known fluorescent dyes reported in the literature so far such as cyanines, squaraines, BODIPYs, coumarins and xanthenes. [3,4] Among them BODIPY dyes represent one of the most prominent class of fluorophores due to their significant chemical and also photophysical characteristics including sharp absorption and emission bands, high quantum yields, excellent molar extinction coefficients and exceptional photostability. Moreover, their rigid structural core offering the ability to easily fine tuning the photophysical properties through simple chemical transformations.[5] On the contrary, xanthene derived chromophores are best known for their excellent photostability, high fluorescence efficiency, remarkable Stoke shifts and structural versatility. [6] The fusion of BODIPY and xanthene fluorescent motifs into a single chromophoric scaffold emerges as a promising strategy for the development of a hybrid fluorophore core with improved optical properties. [7] In this study, we report the synthesis of BODIPY-xanthene hybrid dyes through a styryl linker, via a Knoevenagel condensation between the asymmetrical 5,7-dimethyl-8-phemyl-BODIPY core and the corresponding substituted xanthene aldehydes. Hybridization is a common strategy used in the literature for the synthesis of monostyryl BODIPY analogues, relying mostly on the symmetrical 1,3,5,7-tetramethyl-BODIPY core having two methyl groups containing acidic hydrogen atoms in positions 3 and 5. This approach offers the advantage of an easier synthetic accessibility of this BODIPY core while on the contrary gives lower yields and more laborious purifications due to the formation of 3,5-distyryl byproduct with similar polarity. According to our proposed idea we used the asymmetric 1,3-dimethyl substituted BODIPY gave the final hybrid dyes in pure form maximizing their yields up to 30-50% due to the use of an excess of the corresponding xanthene aldehydes used. Furthermore, our performed UV-Vis and fluorescence spectroscopy studies proved that our newly synthesized fluorescent molecules resulted a remarkable red shift into the NIR region around 720 nm in comparison with the starting BODIPY core (510 nm). Importantly, our analysis revealed that the presence of the oxygen substitution on the xanthene scaffold significantly influences fluorescence intensity, since the methoxy analogue provided higher emission intensity while the presence of the free hydroxy group resulted a notable fluorescence quenching.

## 2. Results and Discussion

### 2.1 Synthesis and structural characterization

In the context of our efforts, we developed a novel synthetic strategy for the synthesis of two NIR BODIPY-Xanthene hybrid dyes. Our proposed synthetic methodology enabling the use of the corresponding xanthene aldehyde precursors **7** and **8** and the asymmetrical BODIPY scaffold **16** through a Knoevenagel condensation. Towards this approach we were able to isolate the monostyryl BODIPY-Xanthene hybrids **1** and **2** in sufficient purities and moderate yields.

### 2.2 Synthesis of xanthene intermediates 7 and 8

The synthesis began with the preparation of the xanthene-derived aldehydes intermediates **7** and **8** as shown in Schemes 1. First, cyclohexanone **3** reacted with phosphorous tribromide (PBr_3_) in dry DMF and to give the bromoenal **4** in 85% yield. [8]

**Scheme 1.**
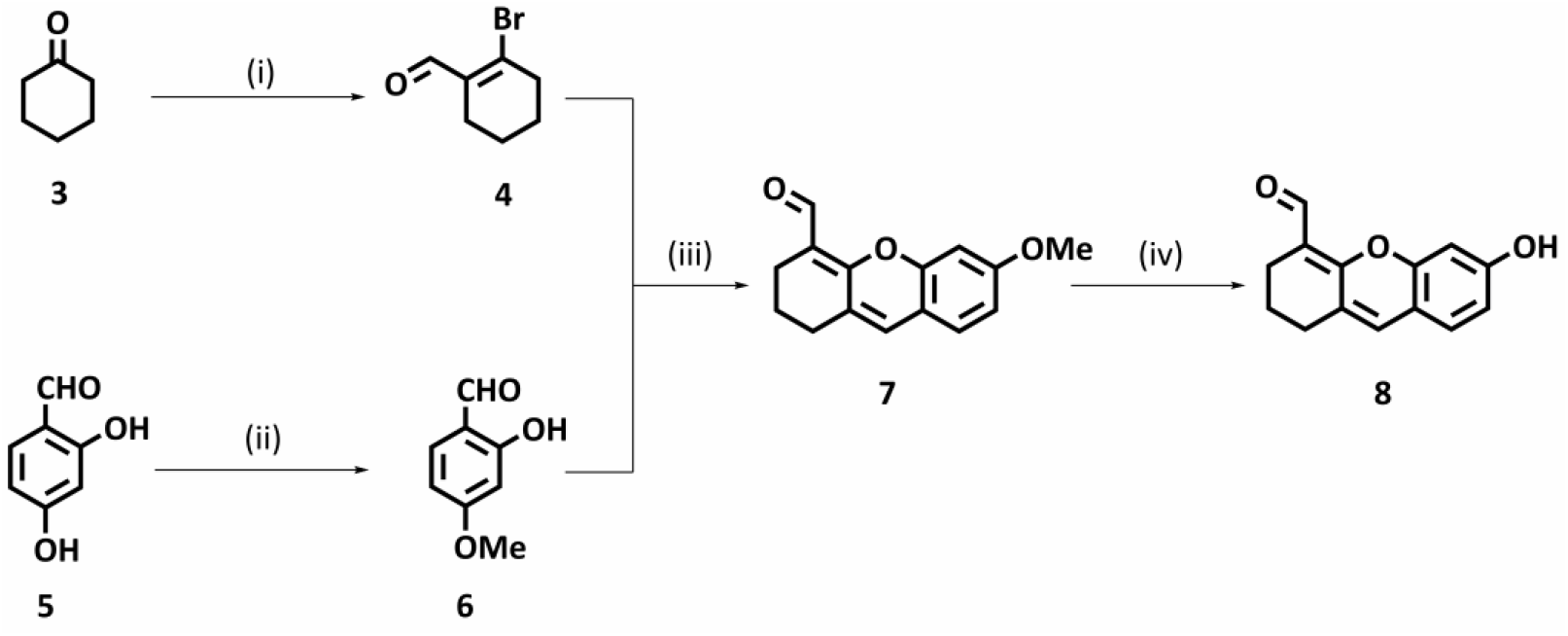
Reagents and conditions: i) PBr_3_, DMF, CHCl_3_, rt, 16 h, 85%; ii) Me_2_SO_4_, K_2_CO_3_, acetone, reflux, 6h, 55%; iii) Cs_2_CO_3_, DMF, rt, 16 h, 75%; iv) BBr_3_, DCM, rt, 16 h, 80%.

Whilst, 2,4-dihydroxybenzaldehyde **5** was methylated using dimemthyl sulfate (Me_2_SO_4_) in the presence of potassium carbonate (K_2_CO_3_) in dry acetone under reflux for 6 h to afford 2-hydroxy-4-methoxybenzaldeyde **6** in 55% yield. [9]

The procedure for the synthesis of xanthene aldehyde intermediate **7** is based on a single step cascade reaction involving, a base-mediated nucleophilic attack of 2-hydroxy-4-methoxybenzaladeyde **6** to the bromocyclohexenecarbaldeyde **4**, followed by a subsequent intramolecular cyclisation under basic conditions, cesium carbonate (Cs_2_CO_3_), to afford xanthene aldehyde intermediate **7** in 75%. [10] The synthesis of the free hydroxy xanthene aldehyde **8** was achieved in 80% yield, after a demethylation reaction of xanthene aldehyde **7** with boron tribromide (BBr_3_) at 0 °C. [10]

### 2.3. Synthesis of asymmetric BODIPY dye 16

The synthesis of the asymmetrical 1,3-dimethyl-8-phenyl BODIPY dye **16** was proceeded, through a four-step synthetic route as outlined in Schemes 2. Towards this direction benzoyl chloride **7** was first reacted with morpholine **10** in dry dichloromethane (DCM) in the presence of triethylamine (TEA) as base, to afford amide **11** in 90% yield.[10] The later amide was then reacted with freshly distilled phosphorous oxychloride (POCl_3_) for 24 h, after which pyrrole **12** was added to the reaction mixture and stirred for another 16 h at room temperature. The resulting intermediate **13** was isolated in 85% yield.[11] Then, treatment of **13** with phosphorous oxychloride (POCl_3_) in dry 1,2-dichloroethane (DCE) for

1.5 h, followed by the addition of commercially available 3,4-dimethylpyrrole **14** and the reaction mixture was refluxed for 2 h to afford dipyrromethene **15** in 85% yield. Finally, the later intermediate **15** was treated with BF_3_.Et_2_O at 0 °C in the presence of TEA in dry DCE, to give the asymmetric BODIPY core **16** in 30% yield.[11]

**Scheme 2.**
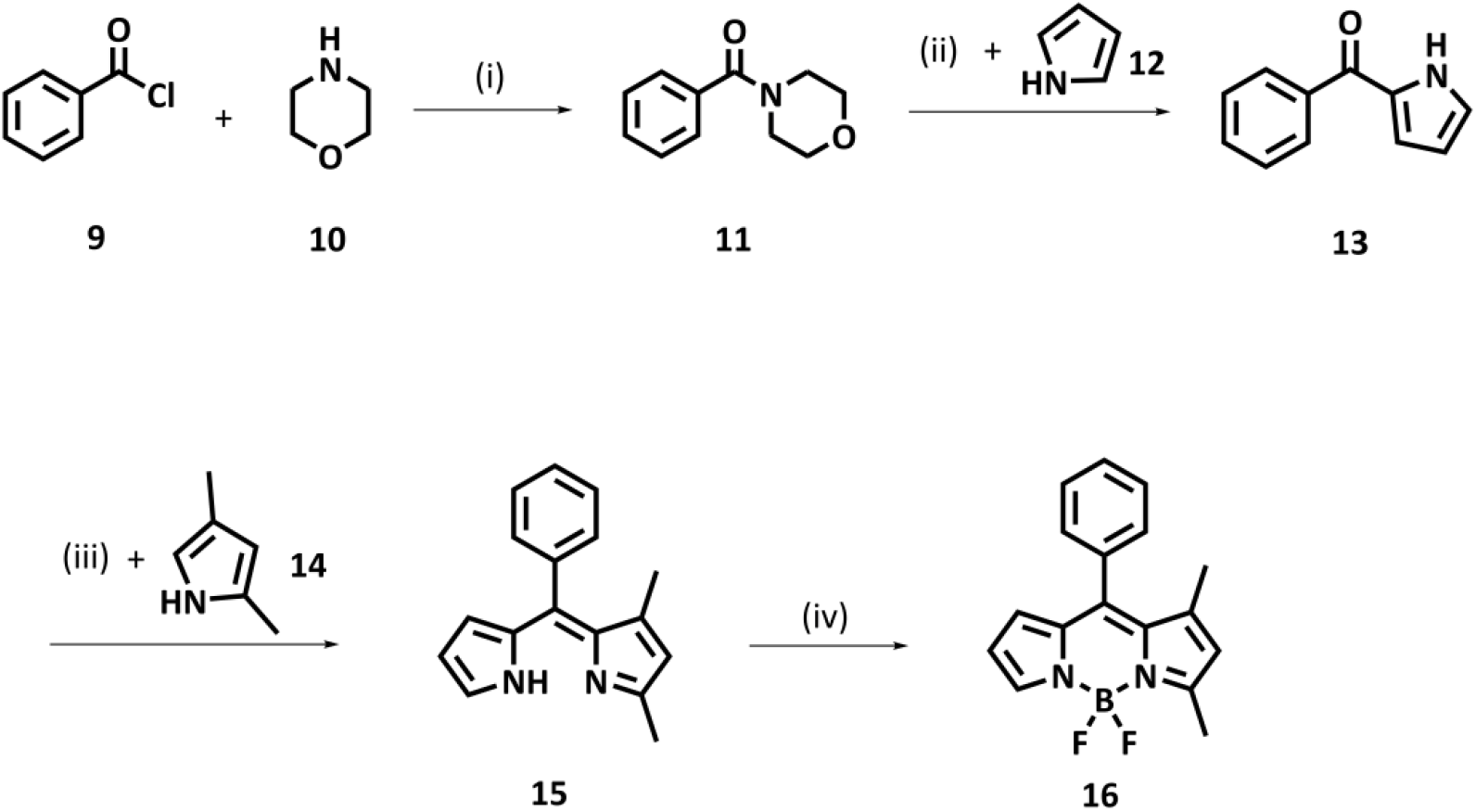
Reagents and conditions: i) TEA, DCM, rt, 16 h, 90%; ii) POCl_3_, + **14**, rt, 48 h, 85%; iii) POCl_3_, DCE, rt, 1.5 h and refluxed 2 h, 85%; iv) TEA, BF_3_.OEt_2_, DCM, rt, 16 h, 30%.

### 2.4. Synthesis of BODIPY-xanthene hybrids dyes 1 and 2

The final synthetic step to obtain monostyryl BODIPY-Xanthene hybrids **1** and **2** (Schemes 3), involved a Knoevenagel condensation between BODIPY **16** and xanthene aldehydes **7** and **8** respectively. The reactions were carried out in the presence of 1:1 mixture of acetic acid and piperidine, at 80 °C for 30 min in the case of hybrid dye **1** and 4h for the hybrid dye **2**. The final hybrid fluorophores **1** and **2** were obtained as blue solids in 50% and 30% yield respectively.

**Scheme 3.**
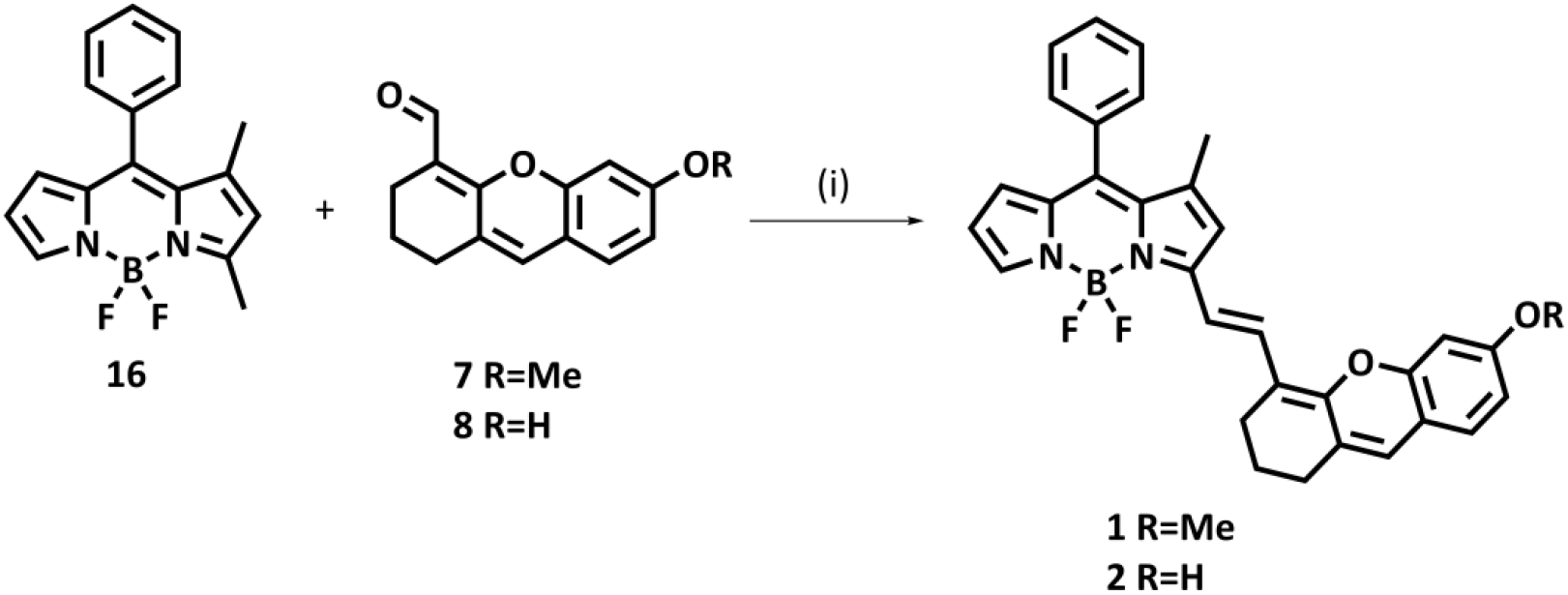
Reagents and conditions: i) acetic acid/piperidine (1:1), DMF, 1.5 h, 80 °C, for **1**: 50%, for **2**: 4h, 30%.

## 3. Optical characterization studies

### 3.1. Synthesis and structural characterization

The optical properties of the synthesized BODIPY–Xanthene hybrids were investigated through absorption and photoluminescence studies. The absorption spectrum of hybrid dye **1** was recorded in THF at various concentrations (ranging from 1.5 to 20 μM), as shown in Figure S1. Hybrid dye **1** exhibits a distinct absorption maximum at 667 nm (Figure 1), the intensity of which shows a linear correlation with the solution concentration (Figure S2). Hybrid dye **1**, bearing a methoxy group on the xanthene core, obeys Beer-Lambert ‘s Law in the range of concentrations that studied in THF, with a high molar extinction coefficient of 7.78 × 10^4^ M^−1^ cm^−1^. (Table 1 and Figure S2)

**Table 1.**
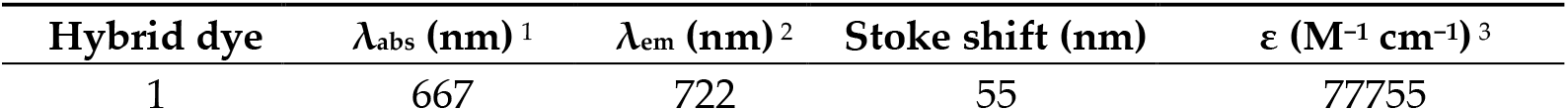
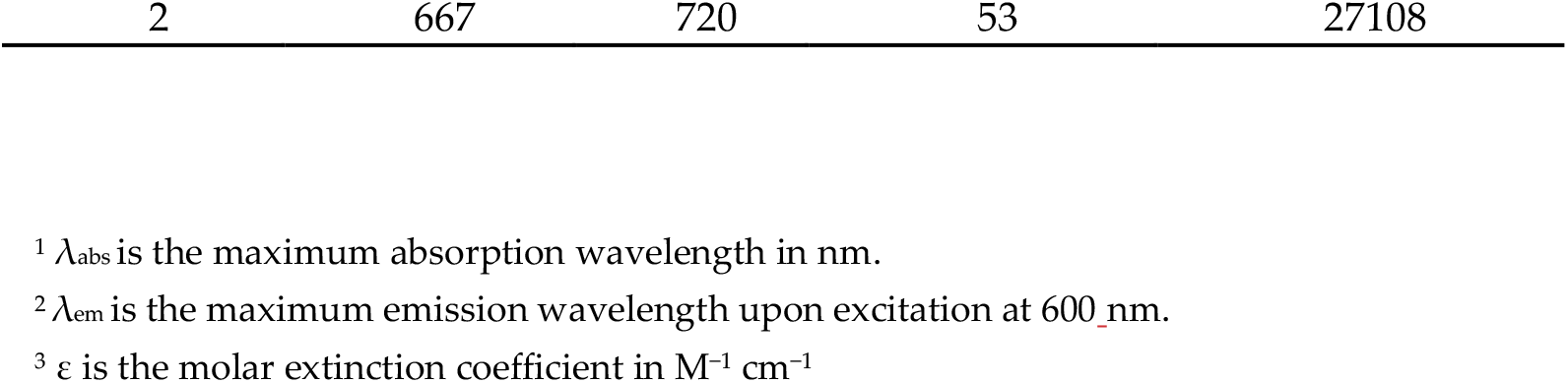
Photophysical properties of BODIPY-Xanthene hybrids in THF.

**Figure 1.**
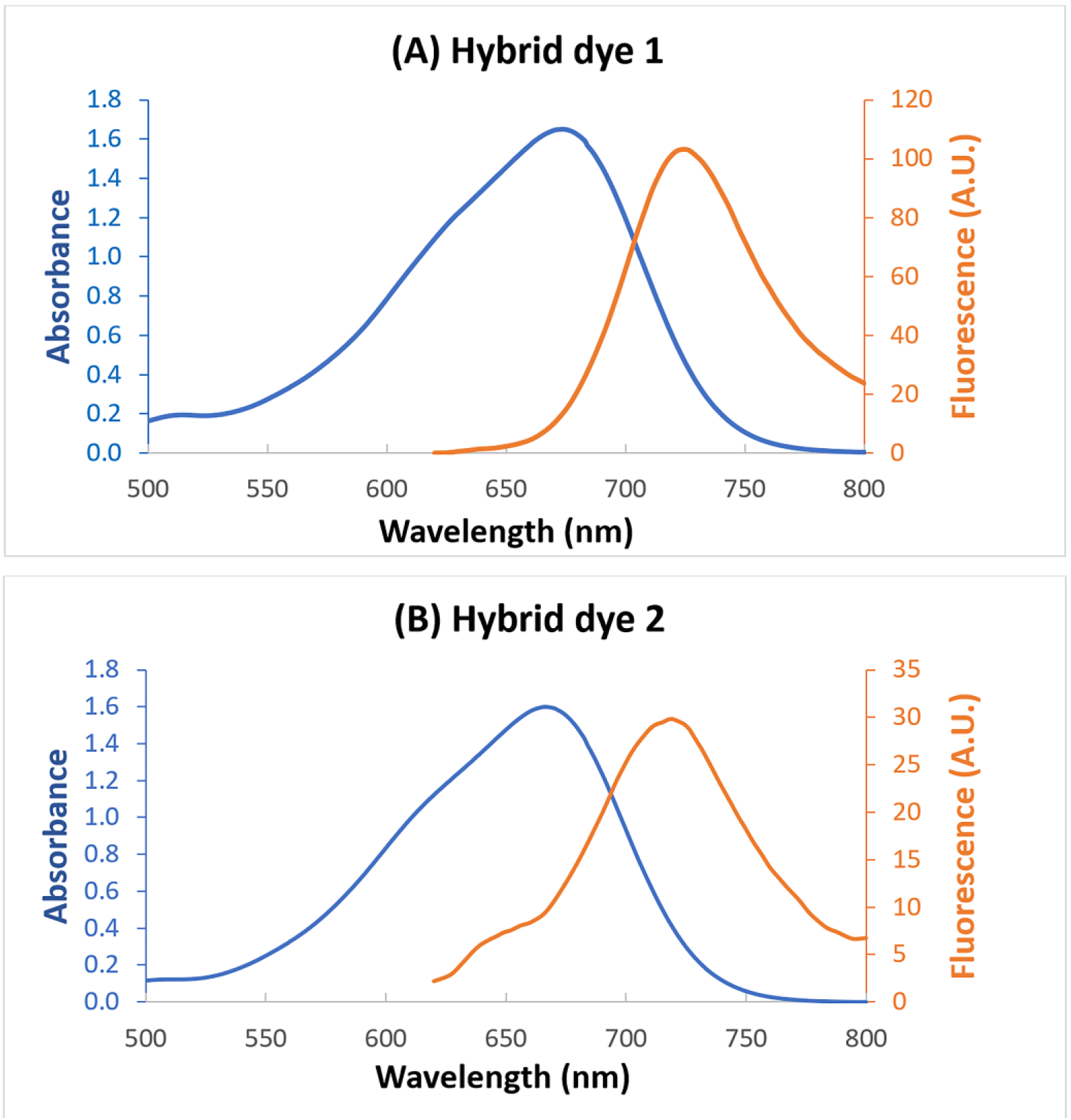
Absorption and emission spectra at 10 μM concentration in THF, (A) of hybrid dye 1 and (B) of hybrid dye 2. Both of them showed maximum absorption at λ_abs_ = 667 nm and emission at λ_em_ = 722 nm and λ_em_ = 720 nm respectively, upon excitation at 600 nm.

The absorption spectrum of hybrid dye **2** was also recorded in THF at concentrations range, from 1.5 to 20 μM (Figure S3). Hybrid dye **2** presents similar absorption spectrum, with maximum absorption also at 667 nm. Based on the results depicted on Figure S3, hybrid dye **2** bearing a free hydroxy group on the xanthene core, also obeys Beer-Lambert‘s Law but with a smaller molar extinction coefficient value of 2.71 × 10^4^ M^−1^ cm^−1^ (Table 1 and Figure S2).

Fluorescence spectra of the two compounds were recorded in THF (10μM) upon excitation at 600 nm. Both hybrids present a remarkable red shift into the NIR region with λ_em_ = 722 nm for hybrid dye **1** (stokes shift 55 nm) and λ_em_ = 720 nm for hybrid dye **2** (stokes shift 53nm) as depicted in Figures 1 and 2.

**Figure 2.**
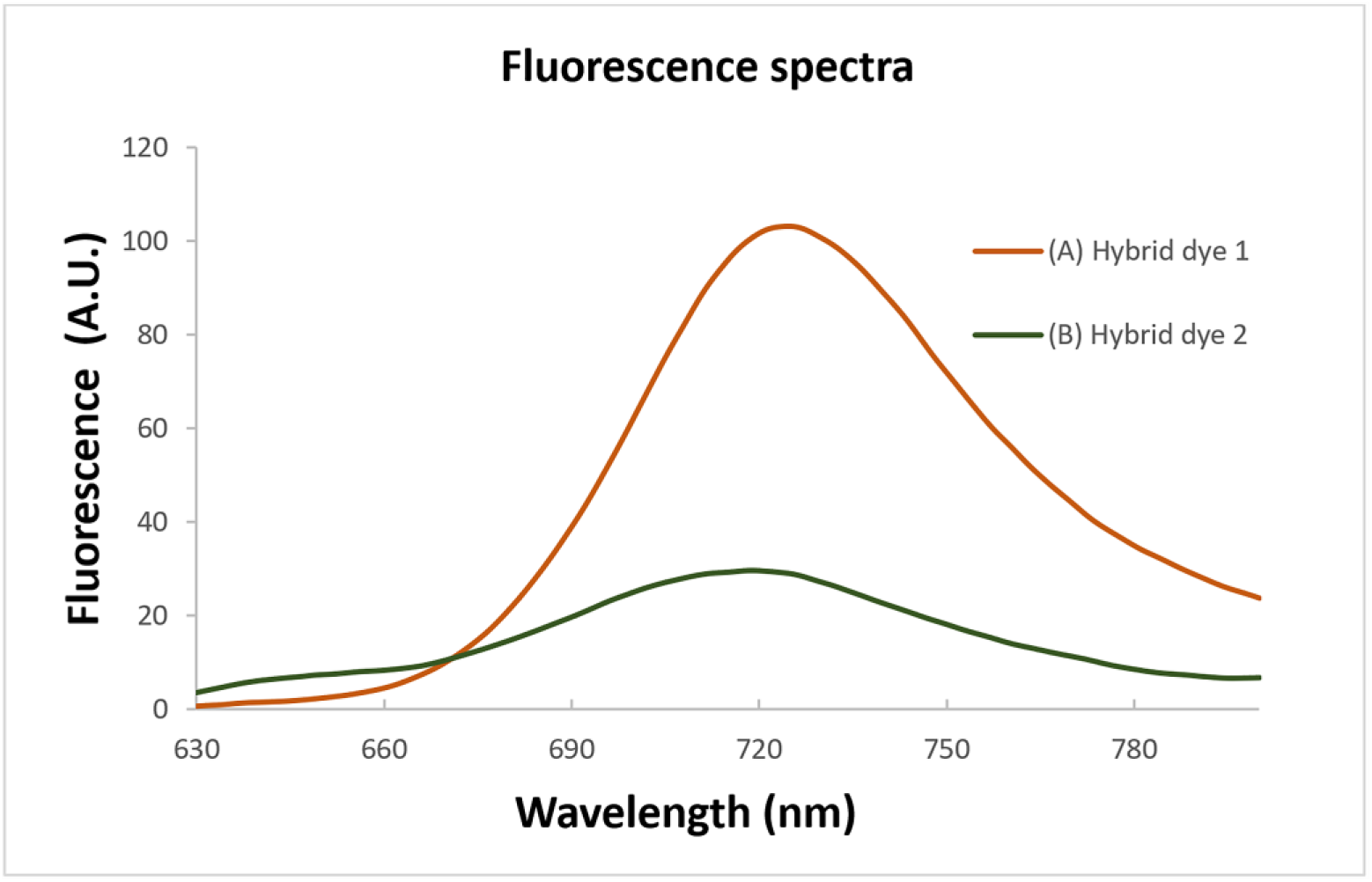
Comparison of the emission spectra at 10 μM concentration in THF, of (A) hybrid dye 1 and (B) hybrid dye 2.

Interestingly, the hybrid dye **1** bearing a methoxy group showed a higher fluorescence intensity than hybrid dye **2** indicating that the free hydroxy group resulted an approximately 70% fluorescence quenching. This effect could be of great importance for future development of optimized NIR fluorescent probes having the OFF/ON fluorescent capability required in optical imaging applications such as real time monitoring of several biological processes or even for the monitoring of drug delivery. (Figure 2)

In order to investigate if the witnessed quenching effect of dye **2** could be due to its potential tautomerism (Schemes 4), we performed DFT calculations and evaluated the dye energy barrier stability in standard conditions (Table 2).

**Table 2.**
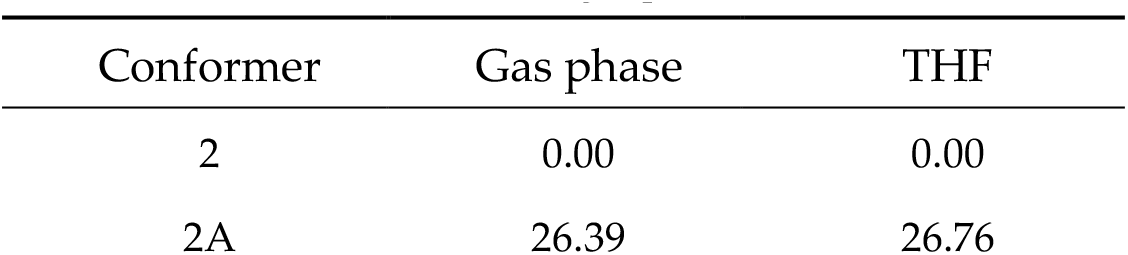
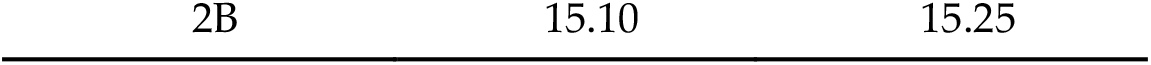
Free Energy difference, ΔG (kcal/mol), of conformers in gas phase and THF.

| Conformer | Gas phase | THF |
| --- | --- | --- |
| 2 | 0.00 | 0.00 |
| 2A | 26.39 | 26.76 |

**Scheme 4.**
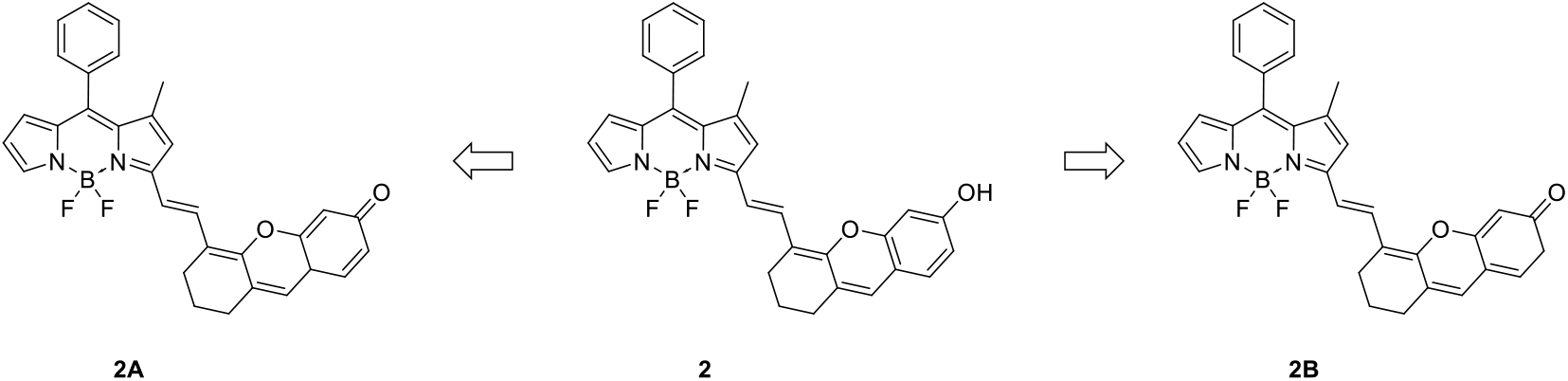
Chemical structures of conformers tested in DFT calculations.

Results suggest that between the tautomer species, 2B showcase a higher existing probability contrarywise to 2A based on ΔG values, with the ∼15 kcal/mol barrier being most likely to interconvert rapidly in solution and coexist in a dynamic equilibrium with 2.

## 4. Materials and Methods

### 4.1. General

Reactions were run in flame-dried glassware under argon or nitrogen atmosphere. All solvents were dried according to standard procedures prior to use. Flash column chromatography (FCC) was performed on silica gel 60 (230–400 mesh, Merck KGaA, Darmstadt, Germany) and thin layer chromatography (TLC) on precoated glass plates 60 F254 (0.2 mm, Merck KGaA, Darmstadt, Germany). Spots were visualized using UV light at 254 and 365 nm and using phosphomolybdic acid stain (PMA, 10% in absolute ethanol). Melting points were determined on an Electrothermal Digital Melting Point Apparatus, Cole-Parmer ET0001/Version 1.0, and were uncorrected. NMR spectra were recorded on Varian spectrometers (Varian Medical Systems, Inc., Palo Alto, CA, USA). ^1^H NMR spectra were recorded at 300 MHz or 600 MHz; ^13^C NMR spectra were recorded at 75 MHz or 150 MHz and were internally referenced to residual solvent peaks. Chemical shifts are reported in δ units, parts per million (ppm). Low resolution mass spectra were recorded on an LC-MSn Fleet mass spectrometer (Thermo Fisher Scientific, Waltham, MA, USA) using MeOH as the solvent. High resolution mass spectrometry (HRMS) spectra were recorded on a UPLC-MSn Orbitrap Velos mass spectrometer (Thermo Fisher Scientific, Waltham, MA, USA). UV-Vis absorption spectra were recorded on a PerkinElmer Lambda 35 UV/VIS spectrometer and fluorescence absorbance measurements on a PerkinElmer Ls 45 fluorescence spectrometer.

### 4.2. Synthesis

#### 4.2.1. Synthesis of 2-bromocyclohex-1-enecarbaldehyde 4

Under anhydrous conditions, 7.7 mL phosphorous tribromide (PBr_3_) was added dropwise at a mixture of dry-DMF (6.95 mL) and chloroform (31 mL) at 0 °C. The reaction mixture was stirred for 45 min at 0 °C. Cyclohexanone **3** (3 g, 0.031 mol) was then added dropwise and the resulting mixture was stirred at room temperature for 16 h. Upon completion, the reaction was carefully quenched in an ice bath and extracted twice with DCM. The combined organic layers were dried over anhydrous Na_2_SO_4_, filtered and concetrated under reduced pressure to afford the crude product **4** as a yellowish oil (4.98 g, 85 % yield). The product was used to the next step without further purification or characterization. [8], [12]

#### 4.2.2. Synthesis of 4-methoxy-2-hydroxybenzaldehyde 6

2,4-dihydroxybenzaldehyde **5** (2 g, 14 mmol, 1eq) was dissolved in dry acetone (40 mL) and stirred under nitrogen atmosphere. Then potassium carbonate (K_2_CO_3_, 2 g, 14.5 mmol, 1.1eq) and dimethyl sulfate (Me_2_SO_4_, 1.37 mL, 14.5 mmol, 1 eq) were sequentially added. The reaction mixture was then refluxed for 6 h and the reaction progress was checked by TLC (PE:EtOAc 9:1, *R*f = 0.64). Upon completion of the reaction, the solvent was removed under reduced pressure and the pure product **6** was isloated by column chromatography (solvent system: PE/EtOAc 95:5 to 9:1) as a white solid (55% yield). [9] ^1^H NMR (600 MHz, CDCl_3_): δ 3.86 (s, 3H), 6.43 (d, *J* = 2.3 Hz, 1H), 6.54 (dd, *J* = 8.6, 2.4 Hz, 1H), 7.43 (d, *J* = 8.7 Hz, 1H), 9.72 (s, 1H), 11.49 (s, 1H); mp 44–46 °C, lit: 41–43 °C [9]; ESI-MS (m/z): 153.16 [M+H]^+^.

#### 4.2.3. Synthesis of xanthene derivative 7

To a solution of 2-bromocyclohex-1-enecarbaldehyde **4** (1.70 g, 9 mmol, 1.4 eq) in anhydrous DMF (50 mL), 2-hydroxy-4-methoxy-benzaldehyde **6** (0.980 g, 6.5 mmol, 1 eq) and cesium carbonate (Cs_2_CO_3_, 6.5 g, 19.3 mmol, 3 eq) was subsequently addded and the reaction mixture was stirred under nitrogen atmosphere at room temperature for 16h. Reaction progress was checked by TLC (PE:EtOAc 9:1, *R*f = 0.15). Upon completion of the reaction (complete consumption of the starting material), the mixture was filtered through a short Ceilite pad which was then washed with ethyl acetate (3 × 20 mL). The filtrates were concetrated under reduced pressure to afford a crude solid product. The solid was dissolved in diethylether and washed twice with water. The organic layer was seperated, dried over anhydrous Na_2_SO_4_, filtered and concetrated under reduced pressure. The desired intermediate **7** was purrified by column chromatography on silica gel 60 (solvent system: PE/EtOAc 95:5) as a yellow solid (75% yield). [12] ^1^H NMR (600 MHz, CDCl_3_): δ 1.70–1.74 (m, 2H), 2.45 (t, *J* = 6.0 Hz, 2H), 2.56–2.59 (m, 2H), 3.85 (s, 3H), 6.66–6.68 (m, 3H), 7.08– 7.10 (m, 1H), 10.30 (s, 1H); mp 143–145 °C; ESI-MS (m/z): 243.1562 [M+H]^+^.

#### 4.2.4. Synthesis of xanthene derivative 8

Compound **7** (200 mg, 0.82 mmol, 1 eq) was dissolved in dry DCM (8.5 mL) and stirred at 0 °C under nitrogen atmosphere. Then a solution of 1M boron tribromide (BBr_3_) in DCM (16.6 mL, 16.4 mmol, 20 eq) was added dropwise and the reaction mixture was stirred at room temperature for 16h. The progress of the reaction was checked by TLC (DCM:MeOH 95:5, *R*f= 0.67). Upon completion of the reaction, it was quenched slowly an aqueous saturated solution of NaHCO_3_ at 0 °C. The mixture was extracted with DCM containing a few drops of methanol. The combined organic layers were dried over anhydrous Na_2_SO_4_, filtered and the solvents recovered under reduced pressure to afford compound **8** as a red solid which was dried under high vacuum over P_2_O_5_ for 2–3 h (0.149 g, 80% yield). [10] ^1^H NMR (300 MHz, DMSO-*d*_*6*_): δ 1.58–1.62 (m, 2H), 2.28 (t, *J* = 6.0 Hz, 2H), 6.43–6.46 (m, 2H), 6.89 (s, 1H), 7.07 (d, *J* = 8.4 Hz, 1H), 10.13 (s, 1H); [13] mp 154–156 °C ; ESI-MS (m/z): 229.19 [M+H]^+^.

#### 4.2.5. Synthesis of phenyl(1H-pyrrol-2-yl)methanone 13

Benzoyl chloride **9** (8.4 mL, 0.073 mol, 1 eq) was dissolved in dry DCM (10 mL) at 0 °C under nitrogen atmosphere. Then, 6 mL of Triethylamine (TEA) was added, followed by a dropwise addition of morpholine **10** (9.6 mL, 0.051 mol, 0.7 eq) at 0 °C and the reaction mixture was stirred at room temperature for 16 h. Upon completion of the reaction, checked by TLC (PE:EtOAc 8:2, *R*f =0.7), the mixture was quenched with 2 M aqueous HCl solution and extracted with DCM (3 × 50 mL). The combined organic layers were dried over Na_2_SO_4_, filtered and the solvent evaporated under reduced pressure. The resulting oily residue was triturated with diethyl ether to afford the intermediate compound **11** as a white solid, after filtration and dried under high vacuum over P_2_O_5_ for 2-3 h (3.5 g, 90 % yield). [11] The amide **11** (1.2 g, 6.3 mmol, 1 eq) was dissolved in phosphorus oxychloride (POCl_3_) (1.6mL, 13.5mmol, 2.15eq) and the reaction mixture was stirred at room temperature for 24h under nitrogen atmosphere. Then, a solution of pyrrole 12 (0.71 mL, 8.2 mmol, 1.3 eq) in dry DCM (2.4 mL) was added and the reaction mixture was stirred at room temperature for an additional 24 h. The reaction was carefully quenched using an aqueous saturated solution of Na_2_CO_3_ at 0 °C under stirring for 30 min. The resulting mixture was then extracted with DCM (3 × 30 mL) and the combined organic layers were dried over Na_2_SO_4_, filtered and concentrated under reduced pressure. The desired product **13** was isolated after purification by column chromatography on silica gel 60 (solvent system: PE/EtOAc, 85:15, *R*f = 0.22) as a white solid (0.90 g, 85% yield). ^1^H NMR (300 MHz, CDCl_3_): δ 6.35–6.35 (m, 1H), 6.91–6.89 (m, 1H), 7.19–7.16 (m, 1H), 7.59–7.46 (m, 3H), 7.93–7.90 (m, 2H); [11] mp 80–82 °C, lit:79–80 °C [14]; ESI-MS (m/z): 172.57 [M+H]^+^.

#### 4.2.6. Synthesis of 1,3-dimethyl-8-phenyl-dipyrromethane 15

Intermedate **13** (0.85 g, 4.97 mmol, 1 eq) was dissolved in dry DCM (21 mL) and phosphorus oxychloride (POCl_3_, 1.86 mL, 20 mmol, 4 eq) was added slowly at room temperature and the reaction mixture was stirred for 1.5 h under nitrogen atmosphere. Then, a s solution of 2.4-dimethylpyrrole **14** (0.612 mL, 5.96 mmol, 1.2 eq) in dry DCE (21 mL) was added and the reaction mixture was refluxed for additionally 2 h. The reaction progress was checked by TLC (PE:EtOAc 8:2). Upon completion of the reaction the mixture was quenched by dropwise addition of an aqueous saturated solution of NaHCO_3_ at 0 °C and extracted with DCM (3 × 30 mL). The combined organic layers were dried over Na_2_SO_4_, filtered and concentrated under reduced pressure to afford bodipy **15** as an orange waxy crude material which was then isolated by column chromatography on silica gel 60 (solvent system: PE/EtOAC 95:5, *R*f = 0.3) as a pure red solid (1.05g, 85%). [11] ^1^H NMR (300 MHz, CDCl_3_): δ 1.39 (s, 3H), 2.36 (s, 3H), 6.01 (d, *J* = 3.8 Hz, 1H), 6.10 (s, 1H), 6.18 (dd, *J* = 3.8, 2.3 Hz, *J* = 3.8 Hz, 1H), 7.22 (s, 1H), 7.31-7.39 (m, 2H), 7.41–7.48 (m, 3H); ^13^C NMR (75 MHz, CDCl_3_): δ 15.2, 18.3, 111.1, 120.9, 126.0, 128.1, 128.3, 129.7, 135.5, 137.5, 137.8, 145.5, 147.4, 167.1; mp 94–96 °C; HRMS: m/z calcd for C_17_H_16_N_2_ [M+H]^+^ 249.1386, found m/z 249.1378.

#### 4.2.7. Synthesis of asymmetric BODIPY 16

Dipyrromethane **15** (0.3 g, 1.21 mmol, 1 eq) was dissolved in dry DCM (12 mL), and stirred at 0 °C under nitrogen atmosphere. After 10 min, triethylamine (TEA, 1.21 mL, 8.7 mmol, 7.2 eq) and boron trifluoride etherate complex(BF_3_.Et_2_O) (1.21 mL, 9.8mmol, 8 eq) were added sequentially. The reaction mixture was allow to stirred at room temperature for 16 h. Then, the reaction mixture was quenched by the addition of saturated aqueous solution of NaHCO_3_, and extracted with DCM (3 × 30 mL). The combined organic layers were dried over Na_2_SO_4_, filtered and concetrated under reduced pressure to afford the product **16** which was isolated as a red solid (0.107 g, 30 % yield) by column chromatography on silica gel 60 (solvent system: PE/DCM 85:15, *R*f = 0.21). ^1^H NMR (600 MHz, CDCl_3_): δ 1.53 (s, 3H), 2.63 (s, 3H), 6.13 (s, 1H), 6.36–6.41 (m, 2H), 7.34–7.36 (m,2H), 7.47–7.52 (m, 3H), 7.68 (s, 1H); ^13^C NMR (150 MHz, CDCl_3_): δ 15.1, 15.3, 116.2, 123.3, 123.4, 127,2, 128.6, 128.8, 129.5, 133.7, 134.1, 134.9, 138.8, 143.5, 147.2, 162.2; [15] mp 139–141 °C, lit: 135–137 °C; [15] HRMS (APCI^-^): m/z calcd for C_17_H_15_N_2_BF_2_ [M]^-^ 296.1302, found m/z 295.1296.

#### 4.2.8. General procedure for the synthesis of BODIPY-Xanthene hybrid dyes 1 and 2

1,3-dimethyl-8-phenyl-BODIPY **16** (20 mg, 0.08 mmol, 1 eq) was dissolved in dry DMF (3.5 mL)and stirred under nitrogen atmosphere The corresponding xanthene aldehyde (**7** or **8**, 0.2 mmol, 2.5 eq) was adeed to the reaction mixture. Then glacial acetic acid (2 drops) and piperidine (2 drops) were added and the reaction mixture was stirred at 80 °C until complete consumption of the starting material (checked by TLC). Upon completion of the reaction, water was added and the mixture was extracted with DCM (3 × 30 mL). The combined organic layers were dried over Na_2_SO_4_, filtered and evaporrated under reduced pressure. The desired product **1** or **2** was isolated by column chromatography on silica gel as blue solids.

##### 4.2.8.1. Synthesis of BODIPY-Xanthene hybrid dye 1

The title hybrid compound was synthesized according the above general procudure with a 30minutes reaction time and using xanthene aldehyde **7** (48.4 mg, 0.2 mmol) to afford the final dye **1** as a blue solid after purification by column chromatography on silica gel (20.8 mg, 50 % yield), (solvent system: PE/DCM 95:5, *R*f = 0.15). ^1^H NMR (300 MHz, CDCl_3_): δ 1.58 (s, 3H, overlapped with the water peak), 1.77–1.87 (m, 2H), 2.49–2.66 (m, 2H), 3.84 (s, 3H), 6.25– 6.35 (m, 2H), 6.42 (s, 1H), 6.55–6.67 (m, 2H), 6.79 (s, 1H), 6.95–7.01 (m, 1H), 7.08–7.18 (m, 1H), 7.42 (d, J = 26.3 Hz, 5H), 7.60 (s, 1H), 7.88 (d, J = 15.7 Hz, 1H); ^13^C NMR (75 MHz, CDCl_3_): δ 15.4, 21.0, 25.0, 29.9, 55.8, 100.7, 110.2, 112.7, 114.8, 115.1, 115.8, 120.1, 123.9, 127.0, 128.2, 128.4, 129.1, 129.4, 134.9, 135.6, 135.7, 138.1, 145.4, 152.1, 154.3, 159.9, 161.1; ^19^F NMR (282 MHz, CDCl_3_): δ -142.98 (q, *J*_BF_ = 31.02 Hz); HRMS (APCI^-^): m/z calcd for C_32_H_27_O_2_N_2_BF_2_ [M]^-^ 520.2139, found m/z 520.2147.

##### 4.2.8.2. Synthesis of BODIPY-Xanthene hybrid dye 2

The title hybrid compound was synthesized according the above general procudure with a 4h reaction time and using xanthene aldehyde **8** (49 mg, 0.2 mmol) to afford the final dye **2** as a blue solid after purification by column chromatography on silica gel (12.2 mg, 30 % yield), (solvent system: PE/DCM 3:7, *R*f = 0.15). ^1^H NMR (600 MHz, DMSO-*d*_*6*_): δ 1.55 (s, 3H), 1.74–1.79 (m, 2H), 2.49–2.66 (m, 2H, overlapped with DMSO peak), 2.52–2.54 (m, 2H), 6.17 (d, *J* = 3.6 Hz, 1H), 6.37 (dd, *J* = 2.4, 3.6 Hz, 1H), 6.58 (dd, *J* = 2.4, 8.4 Hz, 1H), 6.79 (d, *J* = 1.8 Hz, 1H), 6.91 (d, *J* = 15.6 Hz, 1H), 7.13 (d, *J* = 8.4 Hz, 1H), 7.26 (s, 1H), 7.44–7.46 (m, 2H), 7.54–7.56 (m, 3H), 8.03 (d, *J* = 15.0 Hz, 1H); ^13^C NMR (150 MHz, DMSO-*d*_*6*_): δ 14.9, 20.5, 24.3, 29.9, 102.3, 111.4, 111.8, 112.9, 113.9, 115.1, 121.0, 122.7, 125.8, 126.2, 127.7, 128.6, 129.1, 129.3, 133.9, 134.2, 134.8, 135.1, 136.4, 136.9, 145.3, 152.9, 153.7, 159.5, 159.6; ^19^F NMR (282 MHz, DMSO-*d*_*6*_): δ -139.80 (m); HRMS (APCI^-^): m/z calcd for C_31_H_24_O_2_N_2_BF_2_ [M-H]^-^ 505.1908, found m/z 505.1904.

### 4.3. DFT calculations

*Ab initio* calculations were performed at the B3LYP/6-31++G(d,p) level of theory [16-18] on a Dell XPS 8960 desktop PC with Windows 11 64-bit OS (Intel i7 CPU processors, RAM 40 GB), using the Orca 6.1.1 software package. [19-25] Respective Orca input files were created by the use of Atomistica Online 2025 server. [26,27] Initially, a geometry optimization step was performed for all conformers and Hessian matrix calculations were carried out to verify that the optimized structures corresponded to a true minimum with no imaginary vibrational frequencies. Subsequently, single-point energy calculations were performed to obtain the final free energy values at standard temperature and pressure (STP) conditions (1 atm and 298.15 K).

### 4.4. Optical properties

UV-Vis absorption spectra were recorded using a 3mL volume of cuvettes (cuvettes volume are 4.5mL) in a PerkinElmer Lambda 35 UV/VIS spectrometer. The wavelength range was 300-800 nm and the scan speed was set at 480 nm min^-1^. Fluorescence spectra were recorded using a 3mL volume of cuvettes (cuvettes volume are 4.5 mL) in the PerkinElmer LS 45 fluorescence spectrometer. All experiments were carried out at room temperature. The samples were freshly prepared just before measurements.

Stock solutions (0.15 mM) of each dye in THF were prepared. These solutions served as the basis for subsequent dilutions in both absorption and emission studies.

To determine the molar extinction coefficients (ε), absorbance at λ = 667 nm was plotted against the concentration for each dye. The resulting graphs demonstrate a linear relationship, in accordance with the Beer–Lambert law, and the molar extinction coefficient was calculated from the slope of each linear fit.

## 5. Conclusion

In summary, we synthesized two novel BODIPY-Xanthene hybrid fluorescent dyes absorb and emit light within the Near-InfraRed region. The new hybrid dyes exhibited significant red-shift with absorption and emission maxima at λ_abs_ = 667 nm and λ_em_ = 720 – 722 nm upon excitation at λ_em_ = 600 nm, and significant Stokes shifts, in comparison with those of BODIPY dyes in general, ranging between 53 and 55 nm. In addition, the molar extinction coefficient was measured for both of the dyes **1** and **2**, and found to be ε = 77755 M^−1^ cm^−1^ and ε = 27108 M^−1^ cm^−1^ respectively. Importantly, we discovered that the nature of the oxygen atom on the aromatic ring of the xanthene moiety affects on the fluorescent intensity and resulted a 70% fluorescent quenching in case of the hybrid dye containing the free hydroxy group. This will be of significant importance for future development of BODIPY-Xanthene fluorescent switchers, having a fluorescent off-on ability, enabling their uses in analytical optical imaging applications offering real time monitoring of several biological processes.

## Supplementary Materials

The following supporting information are available online and contain information for spectral data (i.e. UV-Vis, NMR, HRMS).

## Author Contributions

Conceptualization, K.C.P.; methodology, S.K., E.A., K.C.P.; validation, G.N., S.K., E.A. and K.C.P.; investigation, G.N.; data curation, G.N., S.K., E.A. and K.C.P.; writing—original draft preparation, G.N., K.C.P.; writing—review and editing, S.K., E.A. and K.C.P.; visualization, S.K., K.C.P.; software, S.K.; supervision, K.C.P., E.A.; project administration, K.C.P.; funding acquisition, K.C.P. All authors have read and agreed to the published version of the manuscript.

## Funding

K.C.P. would like to thank General Secretariat for Science and Innovation of Greece (GSRI) for financial support, TAEDR 053734, 3GPV-4INDUSTRY. G.N. would like to thank the Hellenic Foundation for Research and Innovation (HFRI) for providing her a Ph.D. fellowship.

## Data Availability Statement

Not applicable.

## Conflicts of Interest

The authors declare no conflicts of interest.

## Notes

### Competing Interest Statement

The authors have declared no competing interest.

