## Supplementary material for "Synthesis and characterization of NIR fluorescent BODIPY-Xanthene hybrid dyes": Prousis KC-revised version_10042026_SI_final.pdf

List of content

**I. Copies of NMR spectra**
$^1$H NMR spectrum of (**6**) in $CDCl_3$ (600 MHz) 3
$^1$H NMR spectrum of (**7**) in $CDCl_3$ (600 MHz) 3
$^1$H NMR spectrum of (**8**) in DMSO-$d_6$ (300 MHz) 4
$^1$H NMR spectrum of (**13**) in $CDCl_3$ (300 MHz) 4
$^1$H NMR spectrum of (**15**) in $CDCl_3$ (300 MHz) 5
$^{13}$C NMR spectrum of (**15**) in $CDCl_3$ (75 MHz) 5
$^1$H NMR spectrum of (**16**) in $CDCl_3$ (600 MHz) 6
$^{13}$C NMR spectrum of (**16**) in $CDCl_3$ (150 MHz) 6
$^1$H NMR spectrum of (**1**) in $CDCl_3$ (300 MHz) 7
$^{13}$C NMR spectrum of (**1**) in $CDCl_3$ (75 MHz) 7
$^{19}$F NMR spectrum of (**1**) in $CDCl_3$ (282 MHz) 8
$^1$H NMR spectrum of (**2**) in DMSO-$d_6$ (600 MHz) 8
$^{13}$C NMR spectrum of (**2**) in DMSO-$d_6$ (150 MHz) 9
$^{19}$F NMR spectrum of (**2**) in DMSO-$d_6$ (282 MHz) 9
**II. Optical studies** 10
Figure S1: Absorption spectra of Hybrid 1 in different concentrations in THF 10
Table S1: Absorbance values of Hybrid 1 at $\lambda_{max}$=667nm,
in different concentrations in THF 10
Figure S2: The linear plot between absorbance values and concentrations,
at $\lambda_{max}$=667nm, ε= 77755cm$^{-1}$M$^{-1}$, $R^2$=0.998 in THF 10
Figure S3: Absorption spectra of hybrid 2 in different concentrations in THF. 11
Table S2: Absorbance values at $\lambda_{max}$=667nm, in different concentrations 11
Figure S4: The linear plot between absorbance values and concentrations,
at $\lambda_{max}$=667nm, ε= 27108cm$^{-1}$M$^{-1}$, $R^2$=0.994 in THF 12
**III. DFT calculations**
Table S3: Single-point energies of compound 2 and tautomers in Orca scoring units (Hartree)
Table S4: Gibbs free energies of compound 2 and tautomers in Orca scoring units (Hartree) 12
Table S5: Gibbs free energies of compound 2 and tautomers converted to kcal/mol 13
Table S6: Entropy (S) of compound 2 and tautomers 13
Table S7: Enthalpy (H) of compound 2 and tautomers 13
Table S8: Zero-point energy (ZPE) of compound 2 and tautomers 13

**I. Copies of NMR spectra**

$^1$H NMR spectrum of (**6**) in $CDCl_3$ (600 MHz)

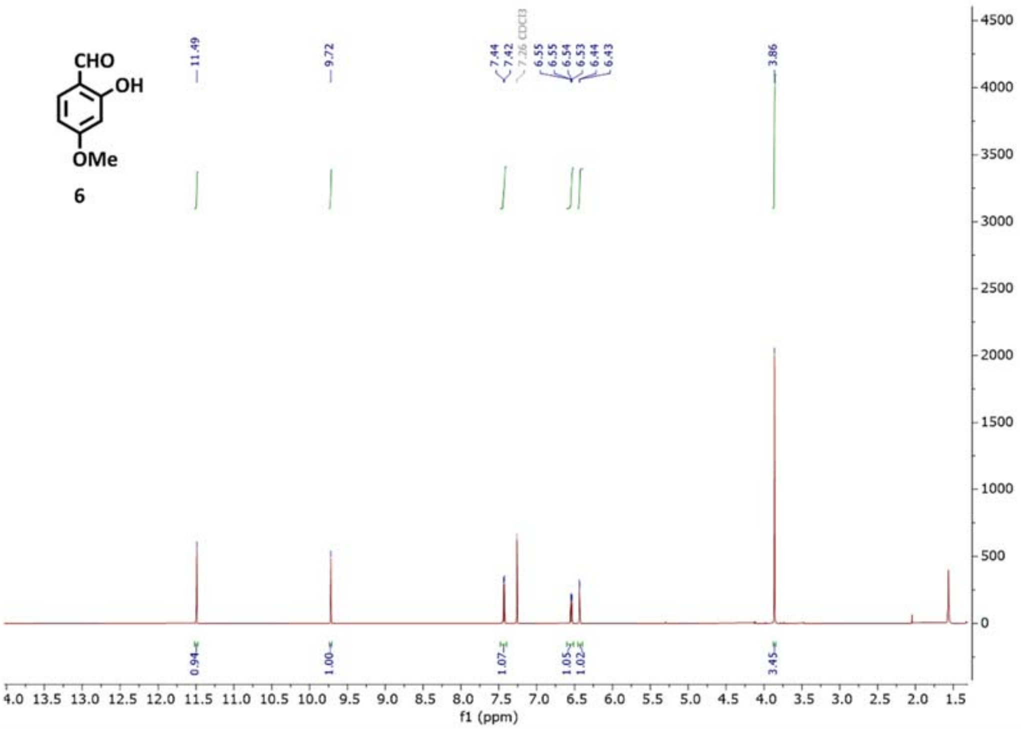

$^1$H NMR spectrum of (**7**) in $CDCl_3$ (600 MHz)

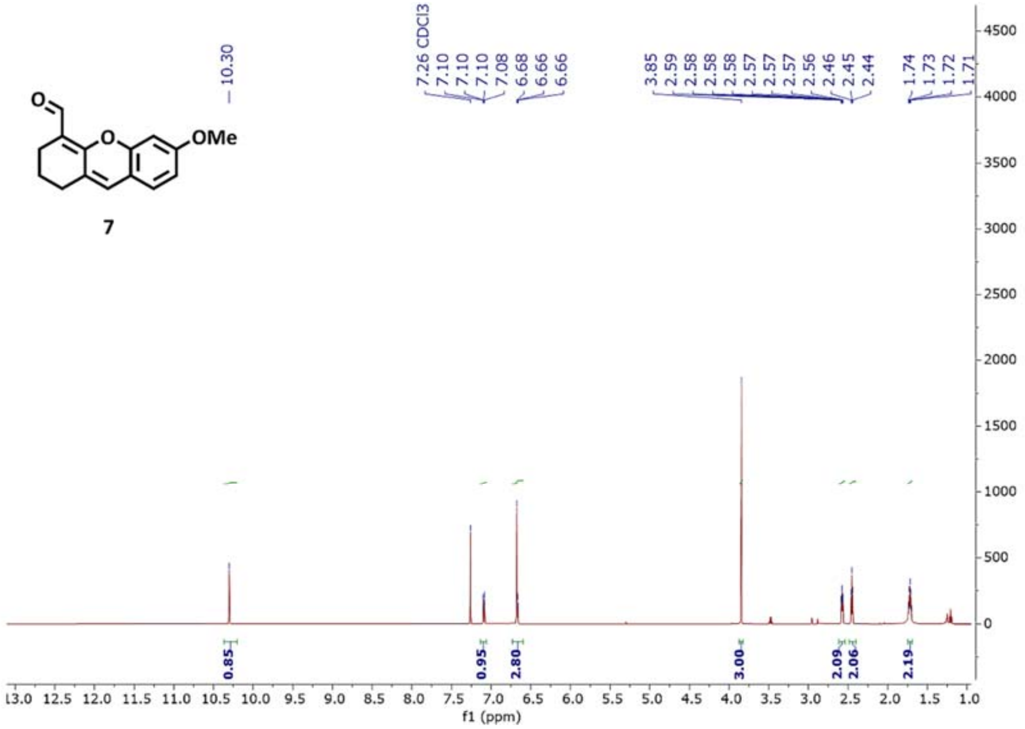

$^1$H NMR spectrum of (**8**) in DMSO-$d_6$ (300 MHz)

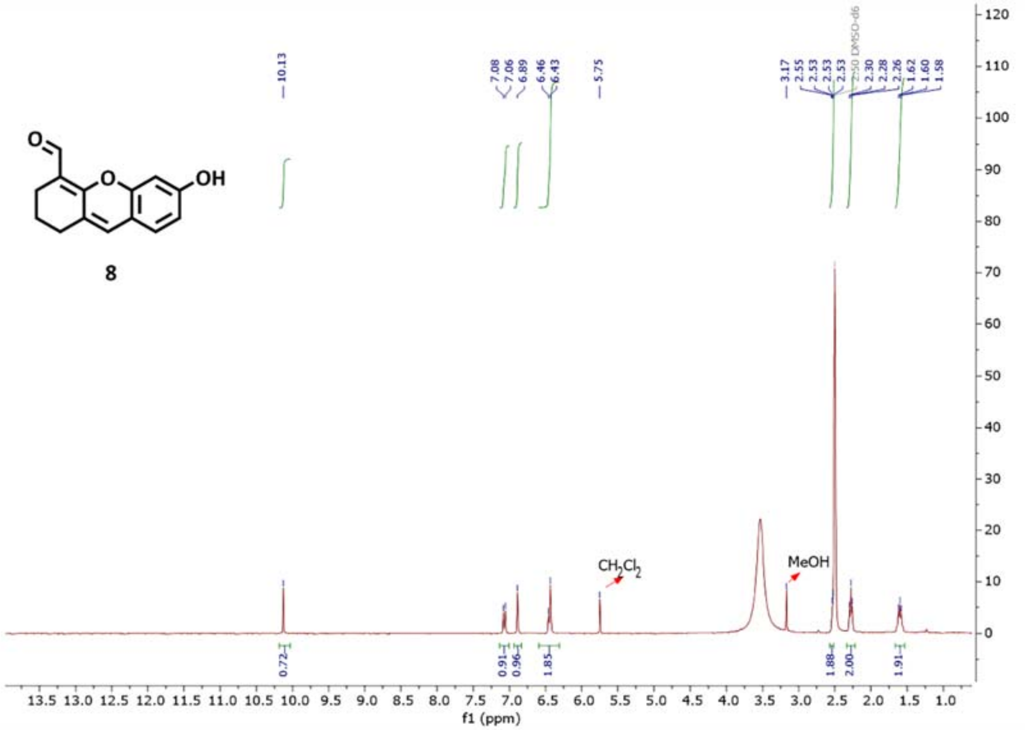

$^1$H NMR spectrum of (**13**) in $CDCl_3$ (300 MHz)

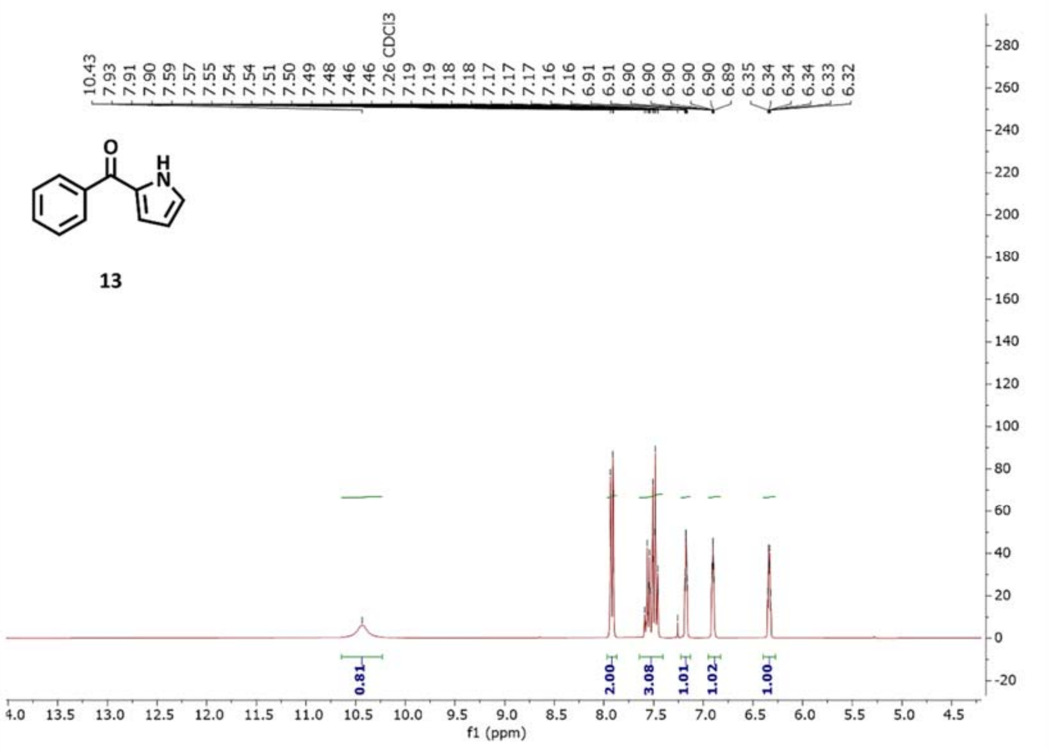

¹H NMR spectrum of (**15**) in $CDCl_3$ (300 MHz)

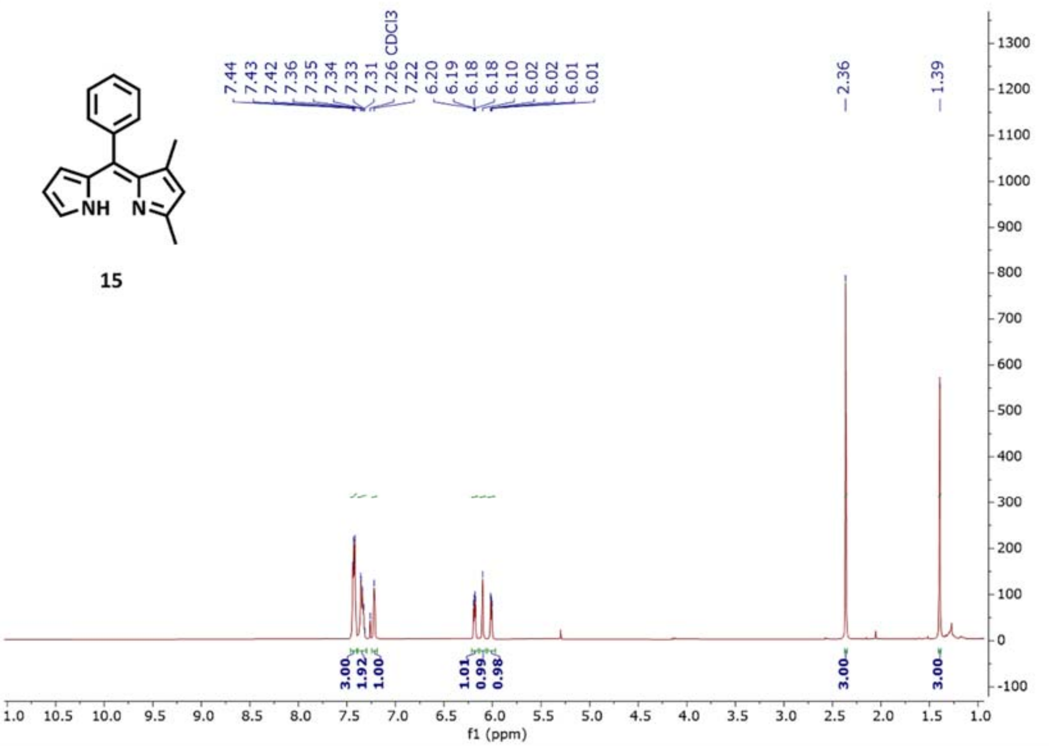

$^{13}C$ NMR spectrum of (**15**) in $CDCl_3$ (75 MHz)

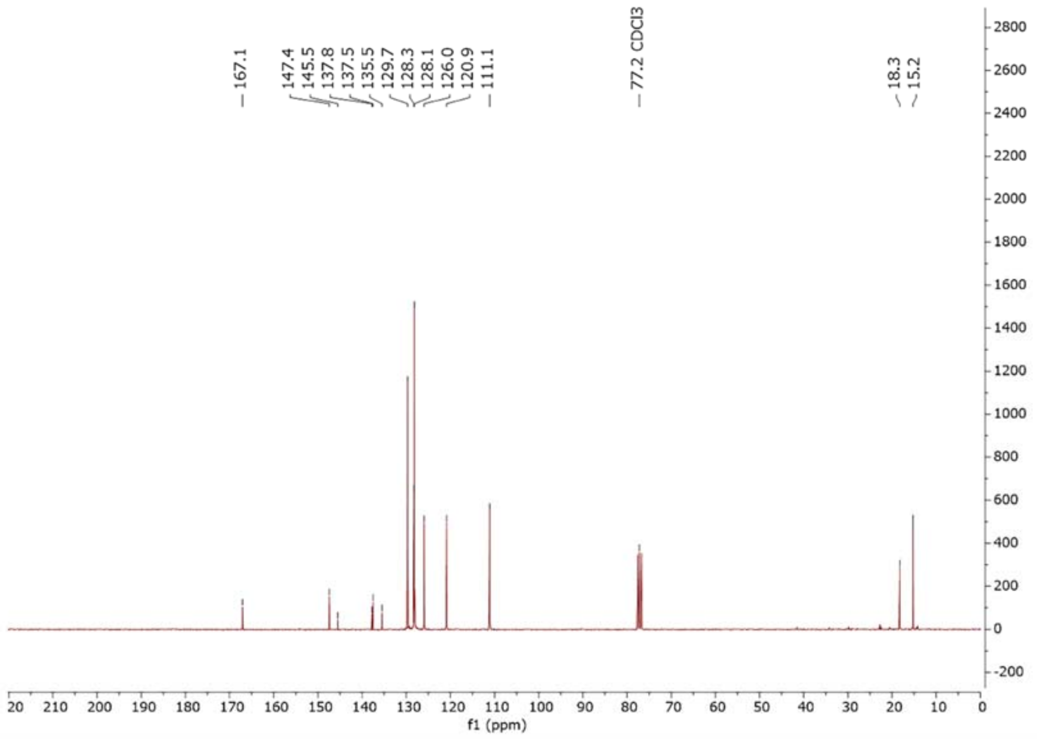


$^{1}H$ NMR spectrum of (**16**) in $CDCl_3$ (600 MHz)

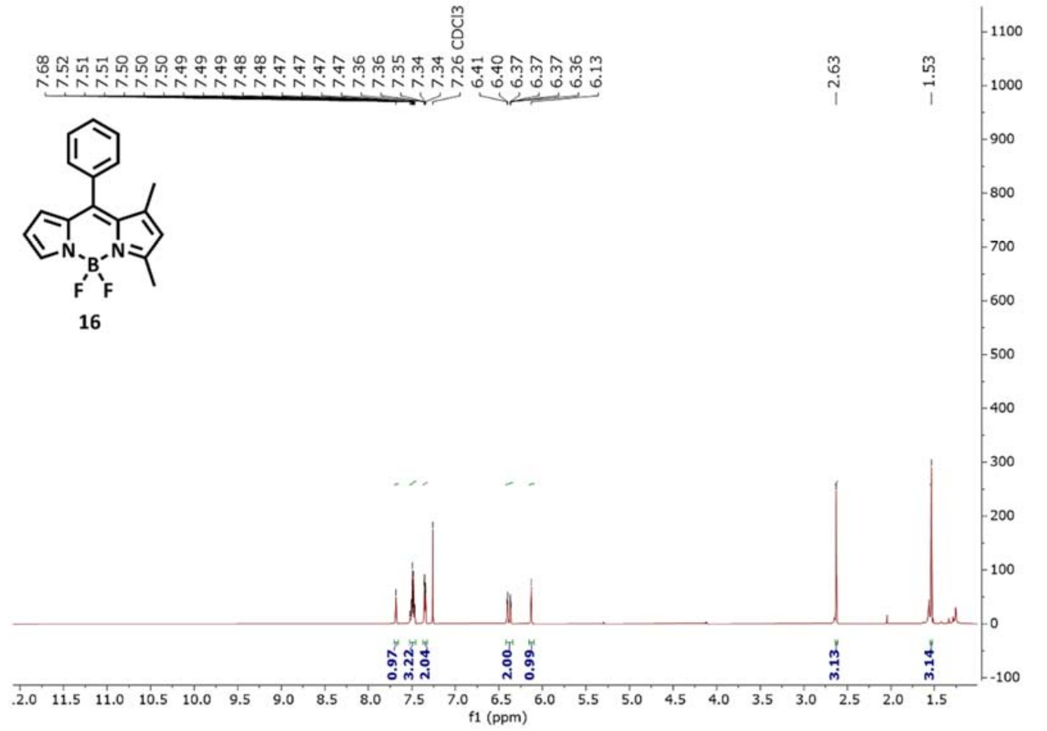

$^{13}C$ NMR spectrum of (**16**) in $CDCl_3$ (150 MHz)

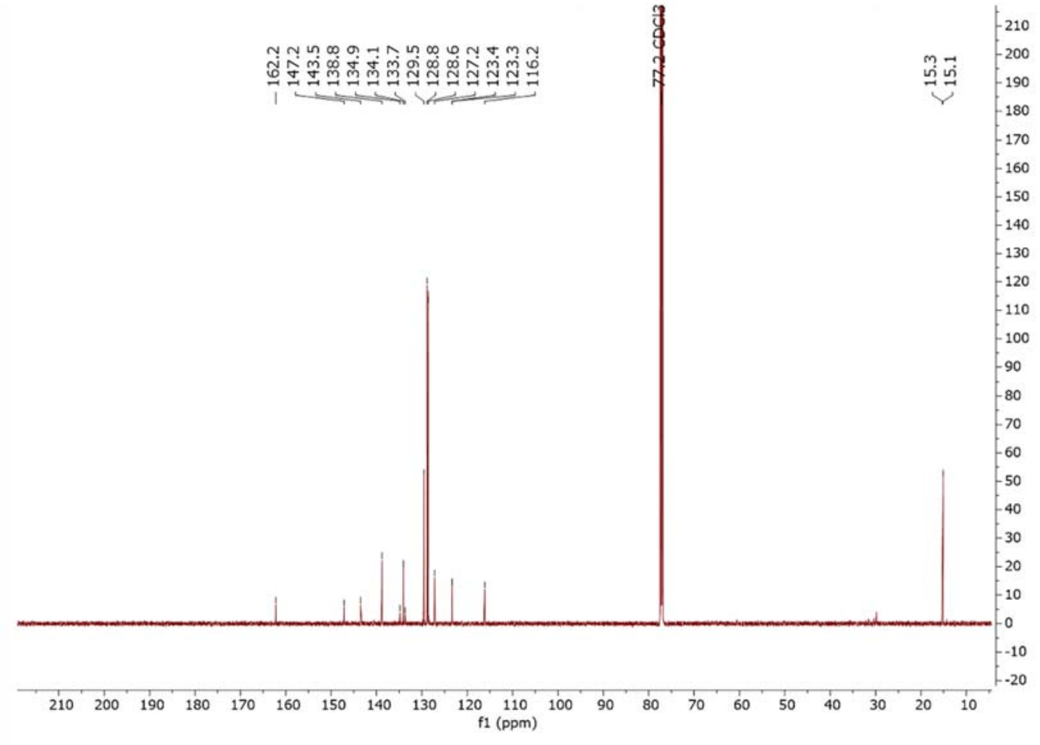

$^{1}H$ NMR spectrum of (**1**) in $CDCl_3$ (300 MHz)

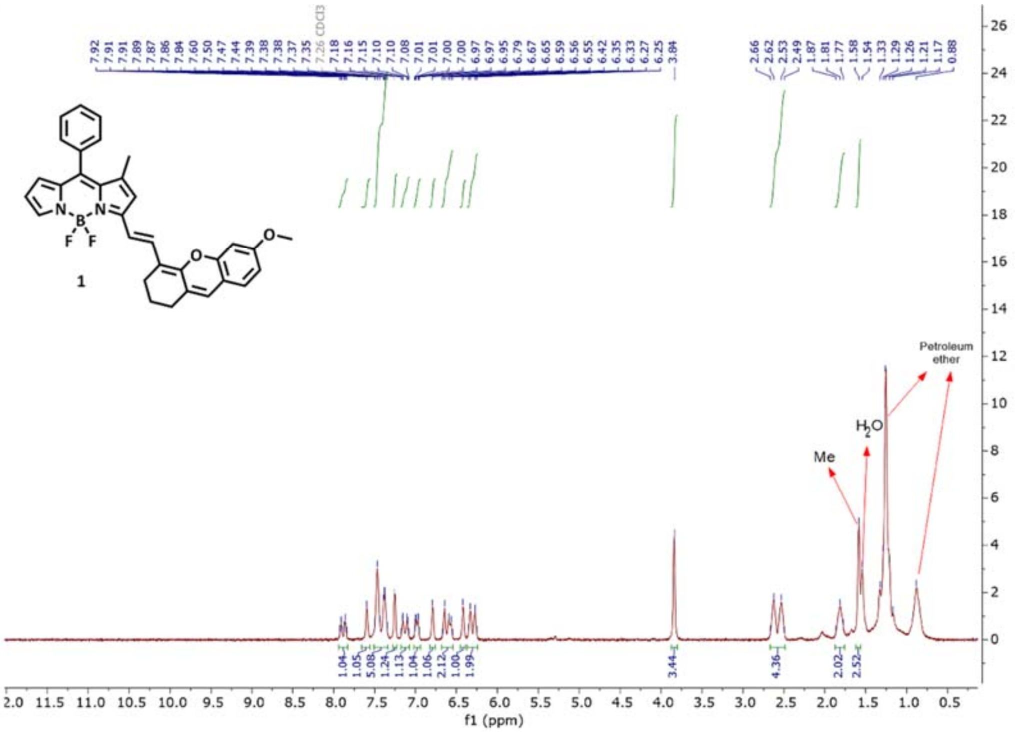

[13] C NMR spectrum of (**1**) in $CDCl_3$ (75 MHz)

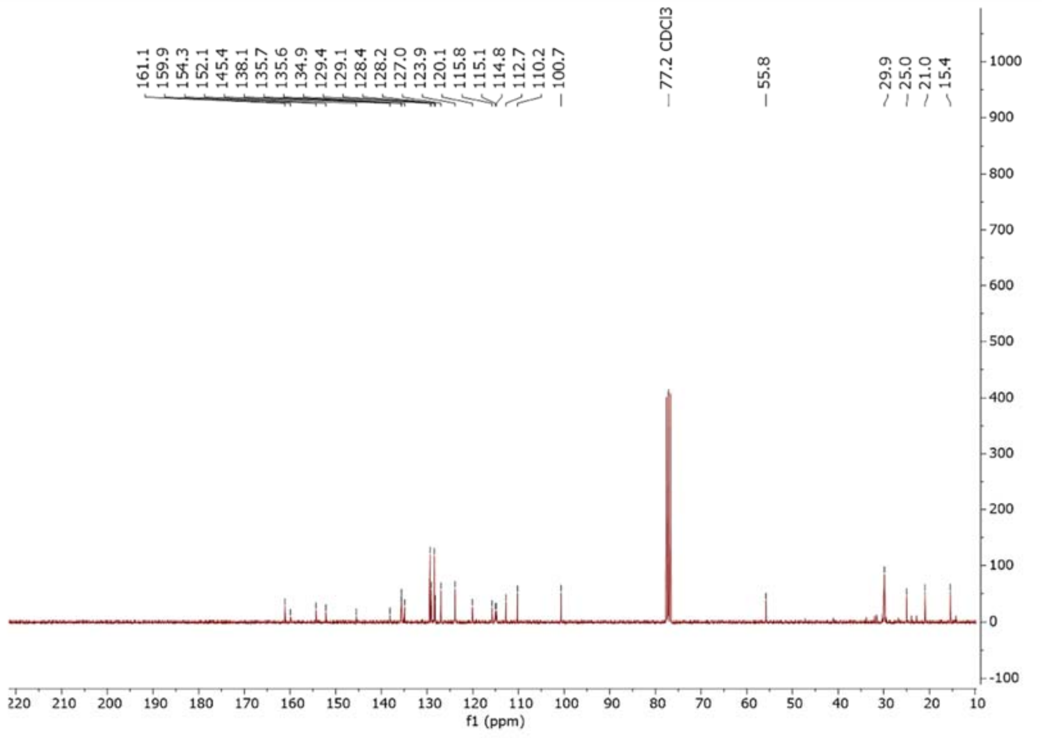


[19] F NMR spectrum of (**1**) in $CDCl_3$ (282 MHz)

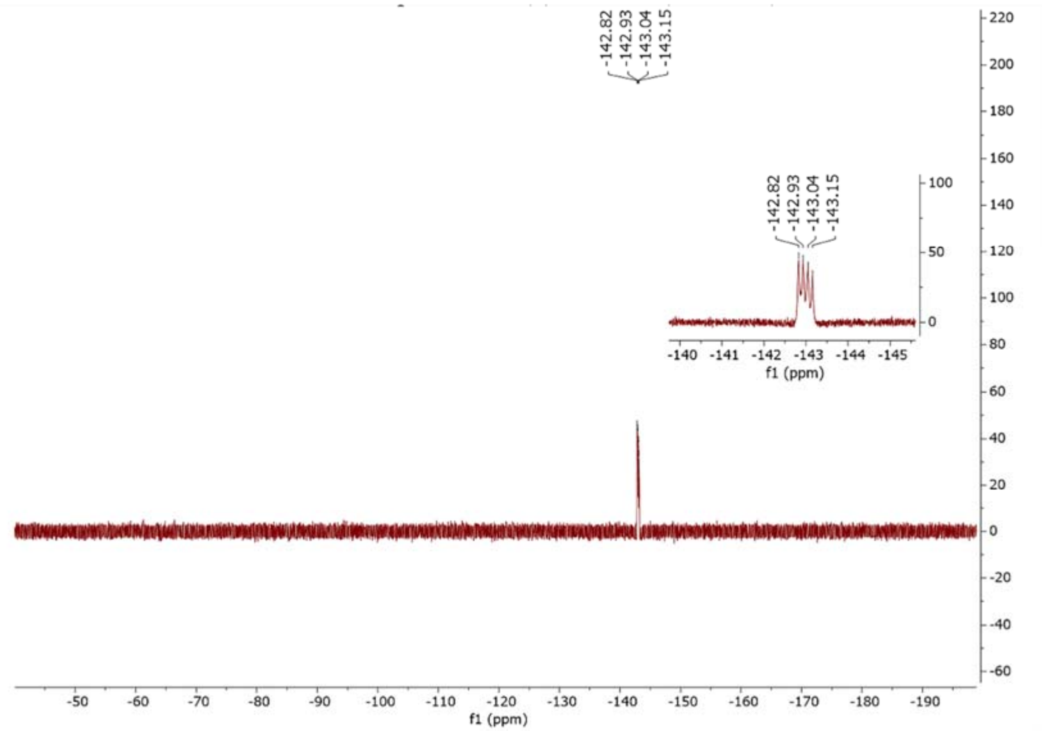

[1]H NMR spectrum of (**2**) in DMSO-$d_6$ (600 MHz)

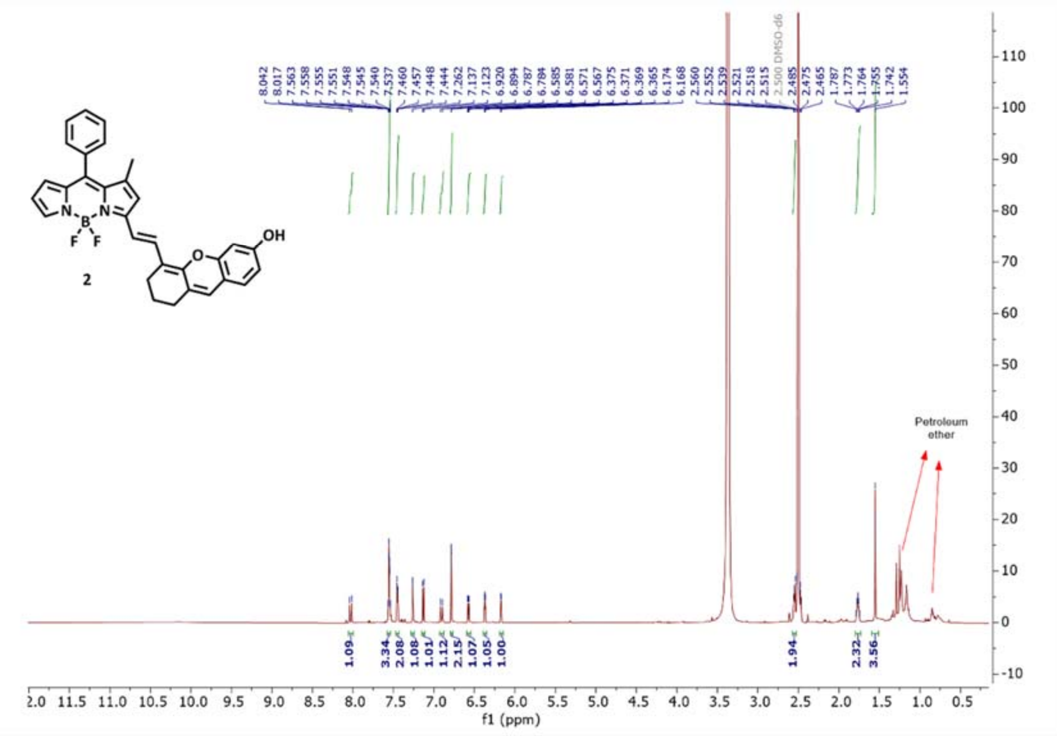

[13]C NMR spectrum of (**2**) in DMSO-$d_6$ (150 MHz)

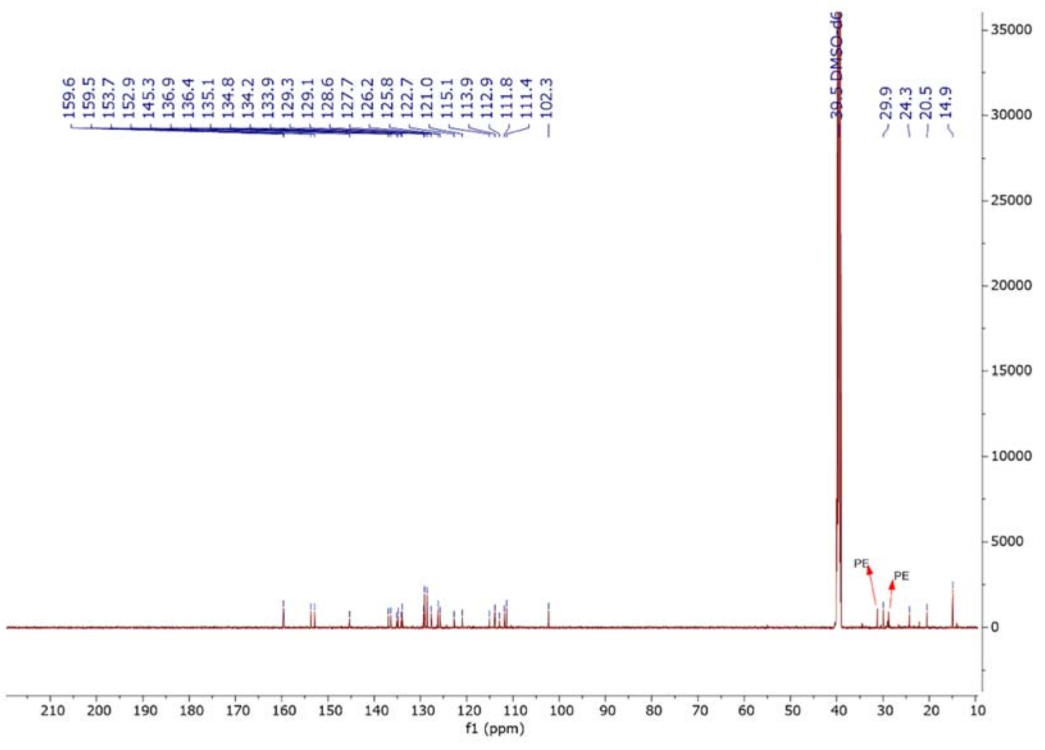

$^{19}F$ NMR spectrum of (**2**) in DMSO-$d_6$ (150 MHz)

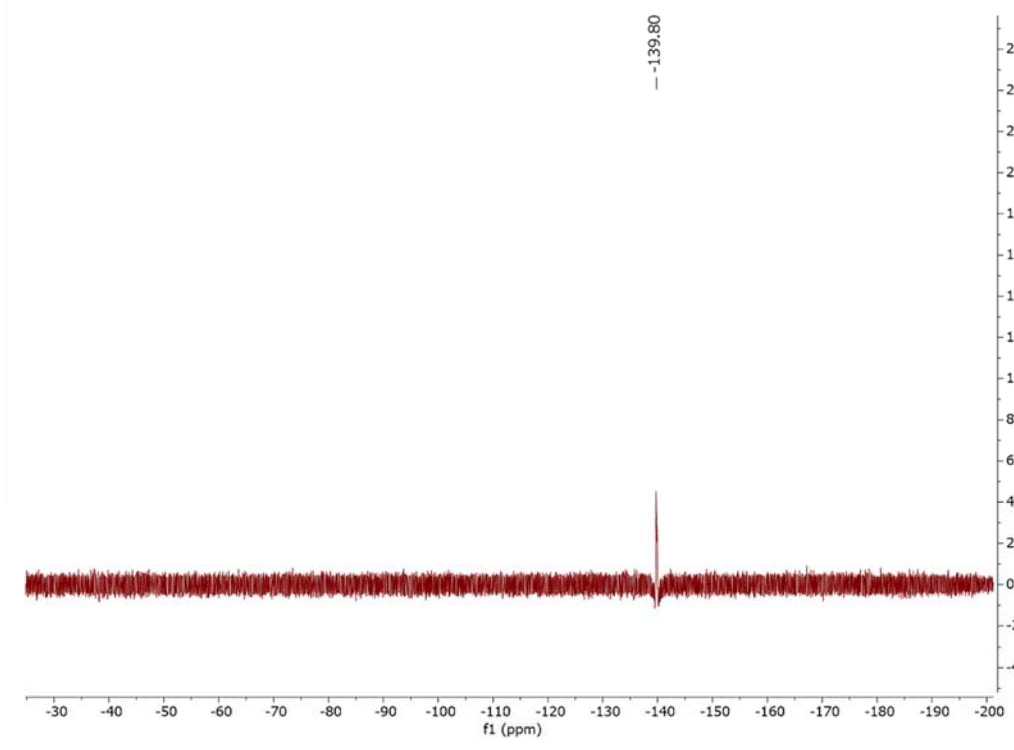


### II. Optical studies

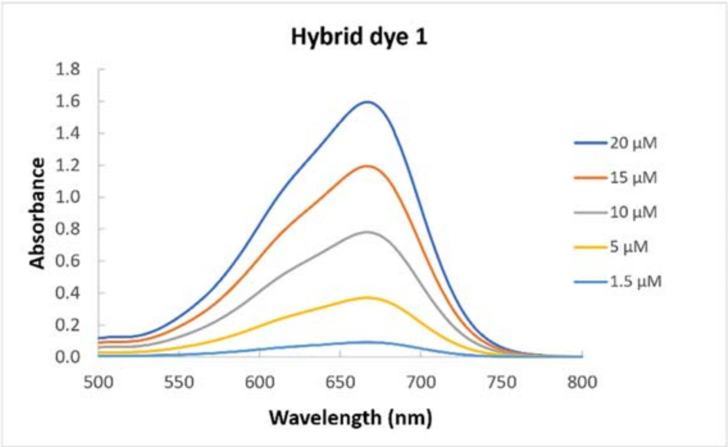


**Figure S1: Absorption spectra of Hybrid 1 in different concentrations in THF.** The absorption spectrum of Hybrid 1 was recorded in THF at various concentrations (from 1.5 to 20 μM).

**Table S1: Absorbance values of Hybrid 1 at $\lambda_{max}$=667nm, in different concentrations in THF.**

| C (μM) | $\lambda_{max}$ (nm) | A |
|---|---|---|
| 1.5 | 667 | 0.091 |

| | | |
|---|---|---|
| *5* | *667* | *0.370* |
| *10* | *667* | *0.780* |
| *15* | *667* | *1.195* |
| *20* | *667* | *1.596* |

The maximum wavelength is recorded at $\lambda_{max}$=667nm, absorbance values in this wavelength were recorded in different concentrations.

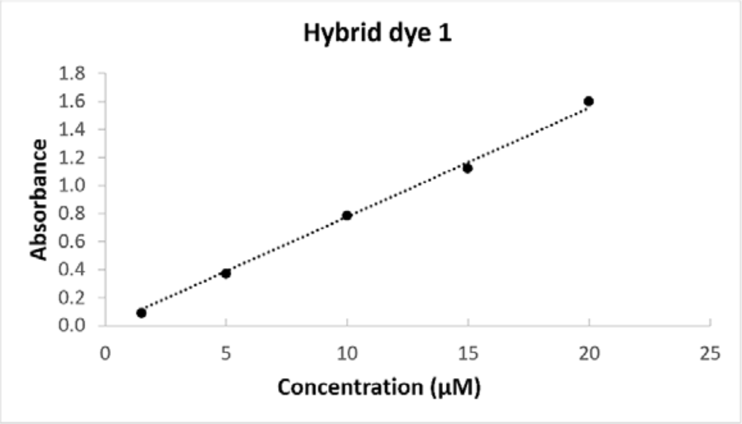


**Figure S2: The linear plot between absorbance values and concentrations, at $\lambda_{max}$=667nm, ε= 77755$cm^{-1}M^{-1}$, $R^2$=0.998 in THF.** According to the Lambert-Beer Law, molar coefficient is ε= 7.78 x $10^4$ $M^{-1}$ $cm^{-1}$, $R^2$=0.9989 in THF.

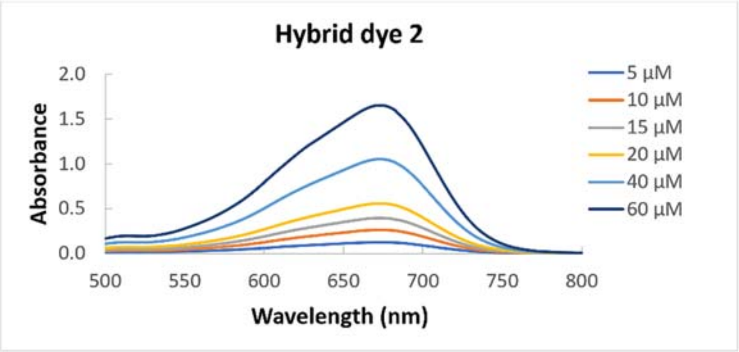


**Figure S3: Absorption spectra of hybrid 2 in different concentrations in THF.** The absorption spectrum of hybrid 2 was recorded in THF at various concentrations (from 5 to 60 μM).

**Table S2: Absorbance values at $\lambda_{max}$=667nm, in different concentrations.**

| C (μM) | $\lambda_{max}$ (nm) | A |
|---|---|---|
| 1.5 | 667 | 0.026 |
| 5 | 667 | 0.125 |
| 10 | 667 | 0.265 |
| 15 | 667 | 0.392 |
| 20 | 667 | 0.555 |
| 40 | 667 | 1.051 |
| 60 | 667 | 1.656 |

The maximum wavelength is recorded at $\lambda_{max}$=667nm, absorbance values in this wavelength were recorded in different concentrations.

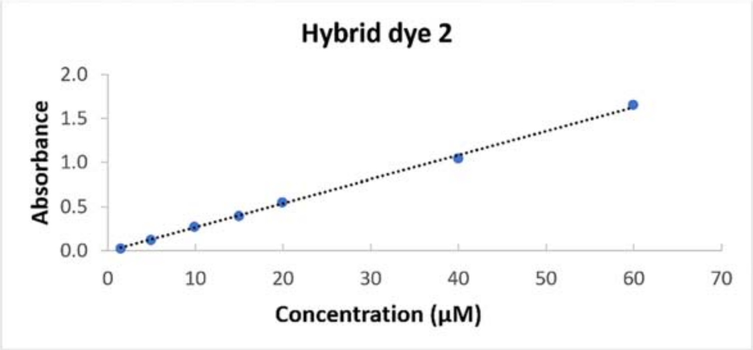


**Figure S4: The linear plot between absorbance values and concentrations, at $\lambda_{max}$=667nm, ε= 27108cm$^{-1}$M$^{-1}$, $R^2$=0.994 in THF.** According to the Lambert-Beer Law, molar coefficient is ε= 27108cm$^{-1}$M$^{-1}$, $R^2$=0.994 in THF.

### III. DFT calculations

**Table S3**: Single-point energies of compound **2** and tautomers in Orca scoring units (Hartree).

| | Compound **2** | Tautomer **2A** | Tautomer **2B** |
|---|---|---|---|
| Single-point energy in gas phase | -1680.1197204898722 | -1680.0775858899049 | -1680.0948023040057 |
| Single-point energy in THF | -1680.1471819546659 | -1680.1051155662253 | -1680.1224522140949 |

**Table S4**: Gibbs free energies of compound **2** and tautomers in Orca scoring units (Hartree).

| | Compound **2** | Tautomer **2A** | Tautomer **2B** |
|---|---|---|---|
| Gibbs free energy in gas phase | -1679.6848945531831 | -1679.6428359397435 | -1679.6608278815513 |
| Gibbs free energy in THF | -1679.7121683960650 | -1679.6695238173552 | -1679.6878592521271 |

**Table S5**: Gibbs free energies of compound **2** and tautomers converted to kcal/mol.

| | Compound **2** | Tautomer **2A** | Tautomer **2B** |
|---|---|---|---|
| Gibbs free energy in gas phase | -1054018.4097959753 | -1053992.0176119416 | -1054003.3077282931 |
| Gibbs free energy in THF | -1054035.5243944316 | -1054008.764511561 | -1054020.2701730477 |

**Table S6**: Entropy (S) of compound **2** and tautomers.

| | Compound **2** | Tautomer **2A** | Tautomer **2B** |
|---|---|---|---|
| Entropy in gas phase | 0.090743567705520112 | 0.090283905258633057 | 0.090752112335431845 |
| Entropy in THF | 0.088817502796620554 | 0.086628471883454072 | 0.087295556330096336 |

**Table S7**: Enthalpy (H) of compound **2** and tautomers.

| | Compound **2** | Tautomer **2A** | Tautomer **2B** |
|---|---|---|---|
| Enthalpy in gas phase | -1679.5941509854777 | -1679.5525520344847 | -1679.5700757692159 |
| Enthalpy in THF | -1679.6233508932683 | -1679.5828953454718 | -1679.6005636957971 |

**Table S8**: Zero-point energy (ZPE) of compound **2** and tautomers.

| | Compound **2** | Tautomer **2A** | Tautomer **2B** |
|---|---|---|---|
| ZPE in gas phase | 0.49316611203638872 | 0.49271241445993164 | 0.49217738309279302 |
| ZPE in THF | 0.49219797060534981 | 0.49150397954035485 | 0.49095192328403026 |
